# Cdc42 mobility and membrane flows regulate fission yeast cell shape and survival

**DOI:** 10.1101/2023.07.21.550042

**Authors:** David M. Rutkowski, Vincent Vincenzetti, Dimitrios Vavylonis, Sophie G. Martin

## Abstract

Local Cdc42 GTPase activation promotes polarized exocytosis, resulting in membrane flows that deplete low-mobility membrane-associated proteins from the growth region. To investigate the self-organizing properties of the Cdc42 secretion-polarization system under membrane flow, we developed a reaction-diffusion particle model. The model includes positive feedback activation of Cdc42, hydrolysis by GTPase-activating proteins (GAPs), and flow-induced displacement by exo/endocytosis. Simulations show how polarization relies on flow-induced depletion of low mobility GAPs. To probe the role of Cdc42 mobility in the fission yeast *Schizosaccharomyces pombe*, we changed its membrane binding properties by replacing its prenylation site with 1, 2 or 3 repeats of the Rit1 C terminal membrane binding domain (ritC), yielding alleles with progressively lower unbinding and diffusion rates. Concordant modelling predictions and experimental observations show that lower Cdc42 mobility results in lower Cdc42 activation level and wider patches. Indeed, while Cdc42-1ritC cells are viable and polarized, Cdc42-2ritC polarize poorly and Cdc42-3ritC is inviable. The model further predicts that GAP depletion increases Cdc42 activity at the expense of loss of polarization. Experiments confirm this prediction, as deletion of Cdc42 GAPs restores viability to Cdc42-3ritC cells. Our combined experimental and modelling studies demonstrate how membrane flows are an integral part of Cdc42-driven pattern formation.

**Significance Statement:** The delivery of new membrane from internal pools at zones of polarized secretion induces in-plane plasma membrane flows that displace slowly mobile membrane-associated proteins from the zone of secretion. However, zones of polarized secretion are themselves specified by the activity of membrane-associated polarity factors, such as the small GTPase Cdc42. Through combined modelling and experimental approaches, this work demonstrates that the fast mobility of the Cdc42 GTPase is critical to allow the establishment and maintenance of a polarity patch, which is reinforced by flow-mediated displacement of a negative regulator.

## Introduction

Polarized secretion, the directional and selective release of molecules from a cell, is critical in many biological processes, including cell communication, tissue development, and immune response. In fungi and other walled cells, it underlies polarized cell growth, as it is necessary for the local delivery of cell wall remodelling and membrane material. Yeast model organisms have been extensively used to understand how the small GTPase Cdc42, which associates with the plasma membrane through a prenyl moiety at its C-terminus, defines the site of polarized growth upon local activation, and dynamically marks the site of polarity.

The Cdc42 polarity system in yeast is believed to represent one of the best examples of biological pattern formation on cell membranes through reaction and diffusion, as initially proposed by Alan Turing (1). This type of pattern formation is driven by non-equilibrium fluxes generated by chemical reactions, combined with spatially varying mobilities (we will use the term “mobility” to indicate both diffusion along the membrane as well as association and dissociation from the membrane). Yeast cells can undergo spontaneous polarization, or symmetry breaking, in which a polarized system arises from a homogeneous one, in absence of a spatial cue. Cdc42 is locally converted to the active, GTP-bound form by the action of a guanine nucleotide exchange factor (GEF). Activation by GEF implements a positive feedback mechanism, where a scaffold protein (Scd2 in *S. pombe*, Bem1 in *S. cerevisiae*) promotes the formation of a complex between Cdc42-GTP and its GEF, leading to activation of neighbouring Cdc42 molecules (2–6). Local Cdc42 activation and amplification leads to one or several zones of Cdc42 activity, as well as the accumulation of Cdc42 at sites of activity. Cdc42 accumulation is thought to be driven by the slower mobility of Cdc42-GTP than Cdc42-GDP at the plasma membrane. In *S. cerevisiae*, thanks in part to extraction of Cdc42-GDP from the plasma membrane by the Guanine nucleotide Dissociation Inhibitor (GDI), this form exchanges with the cytosolic pool more rapidly than Cdc42-GTP, yielding local Cdc42 enrichment (7, 8). In *S. pombe*, the GDI-dependent cytosol-membrane exchange plays a more modest role, but Cdc42-GDP diffuses laterally more rapidly than Cdc42-GTP, so local activation traps Cdc42 (9).

Several mathematical reaction-diffusion models have described Cdc42 polarization mechanistically. A foundational model, which belongs to the prototypical activator–substrate type (10), describes a slowly mobile Cdc42-GTP activator undergoing autocatalytic amplification through the scaffold-mediated feedback described above, fed by the more mobile Cdc42-GDP substrate (11). Subsequent models, all containing critical non-linear amplification of Cdc42-GTP, have for instance explored the role of the GDI (8, 12), of negative feedback (13) and of competition between polarity sites (14) to generate a robust polarity site. However, these models have only implicitly considered the role of GTPase Activating Proteins (GAPs), which are important inhibitory components of the Cdc42 reaction cycle and which are themselves non-uniformly distributed across the cell, to Cdc42 pattern formation. To account for these non-uniform distributions, a model postulated that GAPs switch between fast and slow diffusing states (15).

Prior modelling work has also been inconclusive on the role of the membrane as an evolving element in the Cdc42 system. Consideration of membrane flux is however critical, as the direct outcome of Cdc42 local activity is to promote polarized secretion. Secretory vesicles bring material for cell wall remodelling, which, in walled cells, is essential for local expansion of the cell envelope. Secretory vesicles also bring new plasma membrane, which is largely recycled through endocytosis around the zone of secretion. Cdc42 prenylation occurs in the ER and prenylated proteins are delivered to the plasma membrane by membrane trafficking, but Cdc42 also exchanges between membrane and cytosolic fractions. These physical, non-equilibrium membrane fluxes can in principle provide a driving force for pattern formation, in addition to the classic non-equilibrium fluxes driven by reaction and diffusion.

While polarized delivery of Cdc42 on secretory vesicles had been initially proposed as the main driving force behind Cdc42 polarization (16), this isn’t required for, nor can explain, the accumulation of Cdc42 at sites of polarity (17–19). Replacement of the Cdc42 prenylation site by the RitC peptide (the C-terminal membrane-binding peptide of the mammalian Rit GTPase; (20)), which drives Cdc42 membrane binding directly from the cytosol, also demonstrated that delivery by secretory vesicles is not important for Cdc42 polarization in either organism, as this Cdc42 variant is largely functional and accumulates at sites of activity (9). In some models, membrane delivery was predicted to weaken the polarity site (17, 18) or promote its displacement during pheromone gradient tracking (21).

We recently showed that polarized secretion, coupled with membrane retrieval by endocytosis in a wider zone, leads to the production of in-plane bulk membrane flows away from the zone of membrane insertion (22) (Figure S1). These flows have important consequences on the distribution of membrane-associated proteins, depending on their mobility. For proteins to remain bound to the membrane during transport via membrane flows with velocity *v* (≈ 3 nm/s for *S. pombe*) over length scale *L* (≈ 3 µm for *S. pombe*) requires *L* < *v*/*k*_off_, where *k*_off_ is the dissociation rate. Diffusion with diffusion coefficient *D* will cover a smaller distance compared to such transport when *D*/*v* < *L*. Thus, membrane-associated proteins whose lateral diffusion and dissociation rates are slow enough to satisfy *D*/*v* < *L* < *v*/*k*_off_ are displaced from the secretion zone, as they are unable to dynamically repopulate the naked membrane at the site of secretion (23). For instance, the fast-exchanging small membrane-associated peptide RitC, which exhibits a uniform distribution at the plasma membrane, becomes depleted from secretion zones upon reduction of its exchange dynamics through tandem trimerization (22) (Figure S1A).

Membrane flows directly influence cell patterning by modulating the distribution of Cdc42 regulators. In the fission yeast, three GAPs promote Cdc42-GTP hydrolysis: Rga4 and Rga6 are depleted from cell poles, at least in part due to membrane flows, and accumulate at cell sides (22, 24–26) (Figure S1A). At cell sides, Rga4 and Rga6 actively antagonize Cdc42 activation (25–28). A third GAP, Rga3, co-localizes with Cdc42-GTP zones, but functions additively to the two others, such that triple mutant cells display excessive Cdc42 activity and zone sizes, leading to round cells (28). In these round cells, by constructing an optogenetically-controlled GAP (optoGAP) that couples to membrane flows only in the light, we showed that flow-induced depletion of the Cdc42 GAP from zones of active secretion is necessary and sufficient to promote cell polarization (22) (Figure S1A). Displacement of the GAP to the edge of the secretion zone leaves a zone permissive for Cdc42 activity and corrals it, promoting the formation of the polarity site, and defining its size and the width of the cell.

Do membrane flows also affect the distribution of Cdc42 GTPase itself, despite rapid turnover of the membrane and protein molecules that constitute this site? In this work, we present a reaction-diffusion particle-based model of Cdc42 subject to exo-/endocytosis-induced membrane flows, which shows that robust polarization relies on flow-induced depletion of a low-mobility GAP. We then experimentally reduce Cdc42 mobility at the membrane and demonstrate, in agreement with model predictions, that 1) Cdc42 reduced mobility correlates with expansion of the active zone, 2) strong mobility reduction is incompatible with viability due to GAP-mediated inactivation. Indeed, GAP depletion increases Cdc42 activity, restoring viability at the expense of loss of polarization. Our combined modelling and experimental studies demonstrate how membrane flows are an integral part of the Cdc42-driven pattern formation.

## Results

### Reaction-diffusion particle-based model of Cdc42 with membrane flows

We developed a particle-based model of Cdc42 patch formation which includes the effects of membrane flow together with a reaction-diffusion mechanism for Cdc42 activation and mobility (Figure 1A-C, Materials and Methods). Representing each protein or protein complex as a particle allows a direct implementation of transport by membrane flow (22) as well as accounting for finite-number concentration fluctuations (29). The chemical reaction scheme (Figure 1A) accounts for Cdc42-GDP and Cdc42-GTP membrane diffusion as well as binding and unbinding to the plasma membrane from a cytoplasmic pool of uniform particle concentration. The Scd1/Scd2 GEF complex associates to the plasma membrane or to Cdc42-GTP on the membrane where it catalyzes the conversion of nearby Cdc42-GDP to Cdc42-GTP. This positive feedback retains the essential components needed for active patch formation following a prior particle model for *S. cerevisiae* (29). Slowly diffusing GAPs that catalyze hydrolysis of nearby Cdc42-GDP are also included as discrete particles. These GAP particles represent GAP proteins displaced by membrane flows, such as the optoGAP described above or the endogenous proteins Rga4 and Rga6, which localize to the cell sides (22, 24–26). Therefore, we labeled these discrete GAP particles as sGAPs (slow GAPs). The effect of Rga3, which localizes at cell tips together with Cdc42-GTP (28) is implicitly included in the rate of hydrolysis *k*_hydro_. We do not attempt to describe the competition for Cdc42 between the two tips of *S. pombe* (30); instead we focus on a simple representation of a single tip where we implemented the reactions on a spherical domain. The number of particles in the simulations is based on measured protein or mRNA concentrations. All rates are assumed uniform across the sphere so patterns can only arise via symmetry breaking.

**Figure 1.**
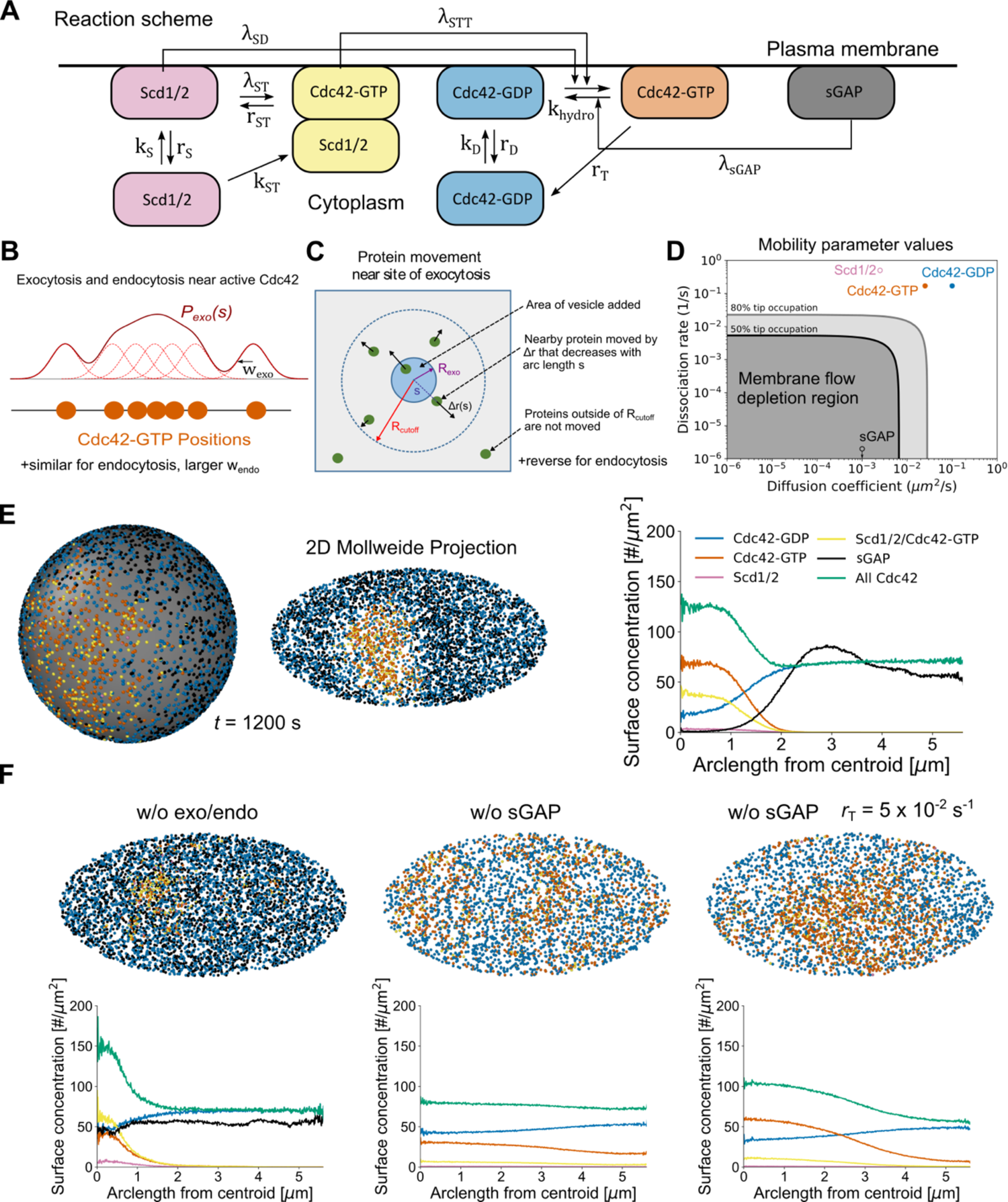
Computational model of Cdc42 polarization in the presence of secretion-driven membrane flows. **A.** Reaction scheme on cell membrane (horizontal line). Cdc42-GTP is formed by two reactions mediated by either Scd1/2 or by Scd1/2/Cdc42-GTP. Boxes not attached to membrane are cytoplasmic. **B.** Exocytosis and endocytosis probability distributions are the convolution of the Cdc42-GTP particle positions and a Gaussian function. **C.** Exocytosis (endocytosis) instantaneously push (pull) particles away from (towards) the site of the event by an amount Δr(s) that depends on their arclength distance from the site of the event. *R_cutoff_* indicates the maximum range of exo/endocytosis. **D.** Reference model mobility parameters. Solid gray and black lines mark boundaries where inert proteins are moderately (80%) or strongly (50%) affected by simulated membranes flows of WT exo/endocytosis centered towards a point of the spherical simulation domain (Figure S1B). Tip occupation ratio is defined as the density of particles near the tip region (< π*R*/4 arclength away from top tip) divided by the density in the remaining area of the sphere (22). **E.** Patch formation occurs for reference case parameters seen from both the 3D sphere, 2D Mollweide projection, and surface concentration profiles from the patch centroid. **F.** Patch formation is weakened at reference parameters by either turning off exo/endocytosis (left) or removing the discrete GAP particle, sGAP (middle). By reducing the dissociation rate of Cdc42-GTP from 0.17/s to 0.05 /s, a wide patch is recovered without sGAP. Snapshots were taken at 1200 s.

The protein particles in the simulation are displaced by vesicle-driven membrane flows, which we model as individual patches of membrane delivered or removed nearby active Cdc42 (Figure 1B). The spatial probability distributions for exocytosis and endocytosis are convolutions of Cdc42-GTP particle positions with a Gaussian of width w_exo_ and w_endo_, which we calculated by comparing tip distributions of CRIB-GFP (marking Cdc42-GTP) to profiles of exocytosis and endocytosis in wild type cells (9, 22). Individual delivery or removal sites were picked from this distribution with a rate matching the corresponding rates of endocytosis and exocytosis at cell tips (22). Particles nearby sites of exocytosis (endocytosis) were moved radially inwards (outwards) and instantaneously by Δr(s), a decreasing function of arc-length distance *s* to the site (Figure 1C). The displacement Δr(s) is based on a model accounting for area conservation of a tensed membrane and is parametrized by two coefficients of order unity describing the fraction of mobile membrane and ratio of lipid to protein velocity (22). The net rate of membrane delivered by exocytosis exceeds that of endocytosis due to net cell growth, which we don’t explicitly simulate (our simulations can be thought to correspond to the frame of reference of a growing tip).

As a result of membrane flows, particles with slow dissociation rate and diffusion coefficients lying within a depletion region are displaced away from a cell tip when using stationary polarized wild type profiles for endocytosis and exocytosis (Figure 1D, S1B) (22). The model assumes that Scd1/2 rate constants lie well outside the depletion region, similar to fast kinetics of the Bem1 complex in budding yeast (29), while sGAP lies within the depletion region (Figure 1D). The dissociation rates and membrane diffusion coefficients of Cdc42-GDP and Cdc42-GTP also lie outside of the depletion region (Figure 1D), as estimated by fitting fluorescence recovery after photobleaching (FRAP) experiments of Cdc42 at cells sides, which we assume is dominated by Cdc42-GDP kinetics, and FRAP recovery at cell tips, which we assume is dominated by Cdc42-GTP (Figure S2; (9)).

Simulations starting with uniformly distributed sGAP, half of all particles (other than sGAP) at random positions on the surface, and reference parameter values (Tables S1, S2) result in spontaneous formation of a single, stably polarized state (Figure 1E, Movie S1). The steady state concentration profiles of individual components (Figure 1E) reproduce the peaks of CRIB-GFP and total Cdc42 observed experimentally in WT cells (9). The steady state endocytosis and exocytosis profiles also match experimental measurements (22) (Figure S3). While the patch of active Cdc42 and GEFs is stable against membrane-driven flow away from the polarization site, sGAP is by contrast cleared away from the active region, similar to the cellular distribution of optoGAP, Rga4 and Rga6 along the cell sides (22, 24–26).

Our model demonstrates how membrane-flow displacement of GAPs is likely an important regulator of pattern formation: removal of vesicle kinetics in the model results in an approximately uniform sGAP distribution and a significantly narrower activation region (Figure 1F). Likewise, deletion of sGAP in the model results in loss of polarization through patch widening (Figure 1F), similar to Cdc42 patch and cell widening in cells lacking one or both of Rga4 and Rga6 (24, 25, 28). We note that the model without sGAP can form patches depending on parameter values and our reference parameter case is on the verge of such symmetry breaking: reducing the dissociation rate of Cdc42-GTP generates a wide patch in the absence of sGAP (Figure 1F).

### Reducing Cdc42 mobility leads to loss of polarity and viability

In previous work, we had shown that the C-terminal membrane-binding region of the mammalian GTPase Rit (ritC) homogeneously decorates the plasma membrane and exchanges rapidly with the cytosolic fraction. By contrast, tandem 2 or 3 ritC repeats lead to reduced detachment rate and progressive depletion from zones of polarized secretion (22). Replacement of Cdc42 prenylation C-terminal CaaX box with 1ritC yields a viable allele, able to promote polarized growth, forming rod-shaped cells (9). To experimentally probe the effect of lowering Cdc42 mobility, we thus constructed Cdc42-2ritC and Cdc42-3ritC variants, predicted to reduce Cdc42 mobility (Figure 2A).

**Figure 2.**
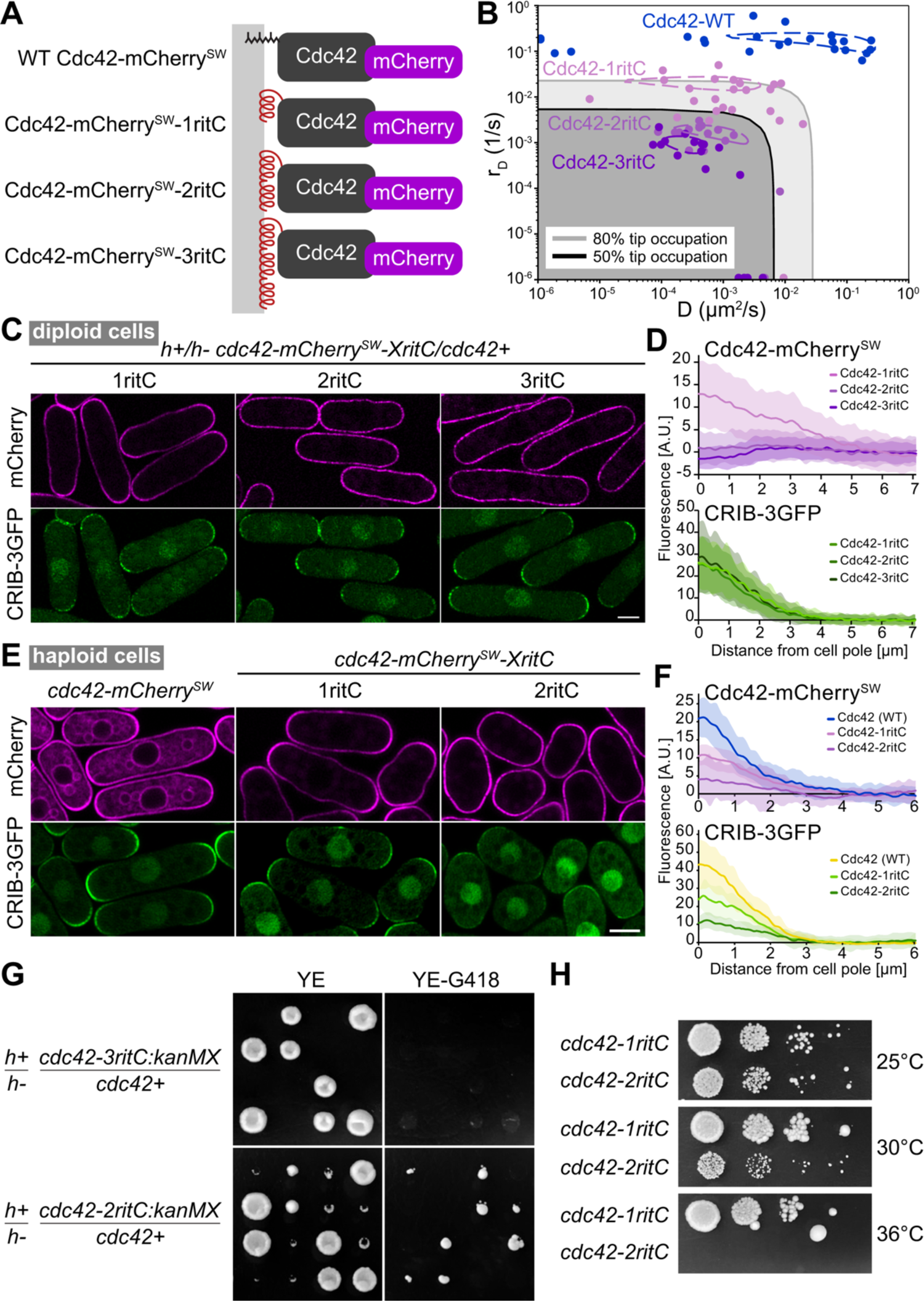
Reduction of Cdc42 mobility at the plasma membrane compromises cell polarity and viability. **A.** Schematic figure of the constructed Cdc42 variants. The prenylation CAAX box is replaced by 1, 2 or 3 tandem copies of RitC (red), which targets Cdc42 peripherally to the plasma membrane (gray). Cdc42 is tagged with mCherry (purple) inserted in a poorly conserved region (9). **B.** D and r_D_ (diffusion and membrane unbinding rates, respectively) of Cdc42-mCherry^SW^ variants, derived from FRAP measurements. The measures were made at cell sides, which captures Cdc42-GDP dynamics. Points are best D and r_D_ fits for individual cells. Dashed contour lines indicate 96% of the maximum R^2^ value of the averaged R^2^ plots for all fits. Gray shaded regions have predicted tip occupation below indicated levels. **C.** Airyscan2 images of Cdc42-mCherry^SW^ variants and CRIB-3GFP (marking Cdc42-GTP) in diploid cells also containing an untagged copy of WT cdc42. **D.** Average Cdc42 and CRIB cortical profiles from cell poles in diploid cells as in (C). Shaded areas show standard deviation. **E.** Averaged airyscan2 images of Cdc42-mCherry^SW^ variants and CRIB-3GFP (marking Cdc42-GTP) in haploid cells. **F.** Average Cdc42 and CRIB cortical profiles from cell poles in diploid cells as in (E). Shaded areas show standard deviation. **G.** Tetrad dissections from indicated diploid strains. Growth on rich non-selective (YE) and *cdc42-mCherry^SW^-XritC*-selective (YE-G418) plates is shown. **H.** Serial 10-fold dilutions of indicated strains on rich medium. Scale bars are 5µm.

We first characterized further the Cdc42-1ritC variant. This and all other Cdc42 alleles described below were also internally tagged with mCherry, shown not to interfere with function (9). We estimated the lateral diffusion and detachment rates of Cdc42-1ritC-GDP through FRAP experiments, which showed that Cdc42-1ritC has somewhat reduced mobility relative to WT Cdc42, but is mobile enough to not be strongly affected by membrane flows (Figure 2B) (22). As previously shown, Cdc42-1ritC accumulated at cell poles (see Figure 2C-F), similar to WT Cdc42 (9). These data establish the baseline against which we can compare Cdc42-2ritC and Cdc42-3ritC variants.

To study the effect of altering Cdc42, we constructed *cdc42-2ritC* and *cdc42-3ritC* alleles in diploid cells, keeping one WT copy of *cdc42*. Estimated lateral diffusion and detachment rates, based on FRAP experiments, indeed showed progressively slower mobilities of Cdc42-1ritC, -2ritC and 3ritC (Figure 2B), similar to the measured mobilities of GFP-tagged 1/2/3ritC (22). In agreement with predictions due to membrane flows (22), reducing Cdc42 mobility led to perturbation of its accumulation at cell poles: in contrast to Cdc42-1ritC, Cdc42-2ritC and Cdc42-3ritC were not enriched at poles, with Cdc42-3ritC cells showing mild local depletion (Figure 2C-D). We note that all diploids exhibited similar distribution of CRIB (Figure 2D), which marks the active Cdc42-GTP pool, and had similar width, indicating that the modified Cdc42 variants do not cause dominant perturbations to polarized cell growth. Thus, tandem dimerization of a membrane anchor can alter Cdc42 distribution, lowering or preventing its accumulation at zones of polarized secretion.

As a test of our model, we performed simulations corresponding to diploid cells by doubling the number of each particle type in the system compared to Figure 1 (Figure S4). To simulate cells with *cdc42-ritC* alleles, half of Cdc42 had diffusion and dissociation constants corresponding to Cdc42-WT and half to Cdc42-1ritC, Cdc42-2ritC, or Cdc42-3ritC, respectively. All cases gave a single polarization site (Figure S4A) and the same (though somewhat stronger) trend of increasing Cdc42 depletion with decreasing Cdc42 mobility (Figure S4B,C, Figure 2D). The model also predicted decreasing Cdc42 activation upon decreasing mobility, which we do not observe in experiments, suggesting that the activity of the WT Cdc42 variant dominates in cells.

To test the functionality of the low-mobility Cdc42 alleles, we sporulated the diploids and dissected tetrads. The *cdc42-3ritC/+* diploid gave rise to only two viable colonies, both lacking the *cdc42-3ritC*-associated selection marker (G418 resistance), indicating that *cdc42-3ritC* is inviable (Figure 2G). The *cdc42-2ritC/+* diploid gave rise to four viable spores, two of which were G418-resistant and slow-growing (Figure 2G). The *cdc42-2ritC* haploid cells were also temperature-sensitive, unable to grow at 36°C (Figure 2H). These cells exhibited aberrant, rounded or stubby cell shape, with increased and variable width, very little CRIB signal at the plasma membrane and almost no CRIB polarization (Figure 2E-F). Similar to the case in diploid cells, Cdc42-2ritC was poorly enriched at cell poles (Figure 2F). In summary, slow-down of Cdc42 mobility decreases Cdc42 activity and accumulation at cell poles, leading to loss of cell polarity and inability to sustain viability.

### Model reproduces experimental observations of Cdc42 mobility reduction

To understand how haploid Cdc42-2ritC cells polarize when Cdc42 itself is severely impacted by membrane flow while haploid Cdc42-3ritC are inviable, we repeated the simulations of Figure 1 but with Cdc42-GDP diffusion and dissociation constants corresponding to those of Cdc42-1ritC, Cdc42-2ritC, and Cdc42-3ritC (Figure 3A). The three Cdc42-ritC constructs have lower Cdc42-GDP diffusion and dissociation constants than WT Cdc42-GTP (Figures 1D, 2B, S2); thus we assumed that the corresponding Cdc42-GTP and Scd1/2-Cdc42-GTP complex parameters are limited by the ritC anchor and set them the same as for Cdc42-GDP.

**Figure 3.**
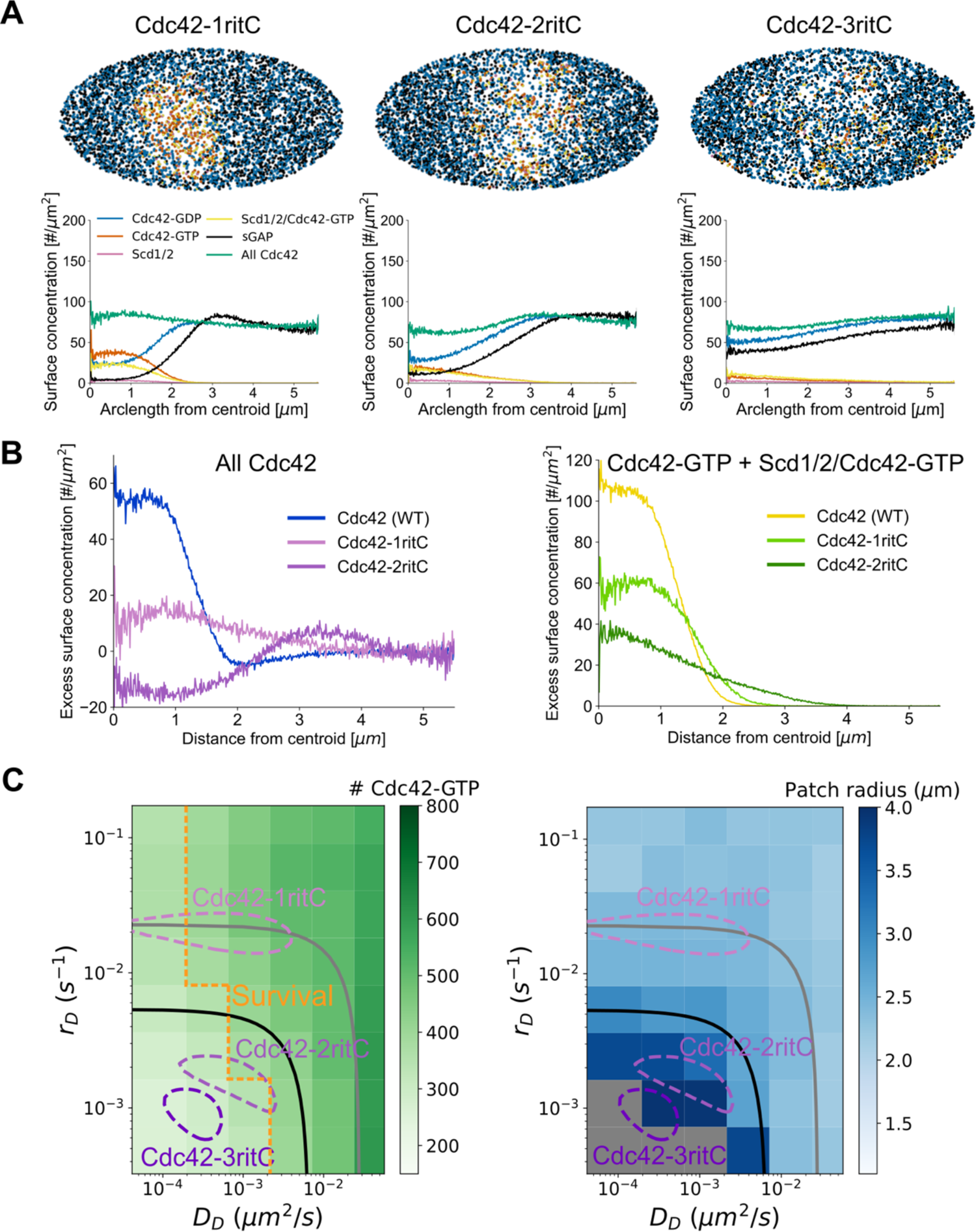
Reduction of Cdc42 mobility in model leads to wider patches and lower Cdc42-GTP levels. **A.** Model results at steady state show that patch width increases as Cdc42 mobility is decreased to parameters for Cdc42-1ritC, Cdc42-2ritC, and Cdc42-3ritC as measured by FRAP. Snapshots taken after 8000 s. **B.** Comparison of total Cdc42 levels and Cdc42-GTP levels (both normalized to side intensity) for Cdc42 (WT), Cdc42-1ritC, and Cdc42-2ritC as a function of distance from Cdc42-GTP patch centroid. **C.** Parameter scan for membrane dissociation, *r_D_*, and diffusion constant, *D_D_*, for Cdc42-GDP, showing total Cdc42-GTP levels (left) and patch radius (right). The Cdc42-1ritC, Cdc42-2ritC and Cdc42-3ritC contour lines and 50% and 80% depletion boundaries are drawn as in Figure 2B. The postulated survival boundary is drawn at approximate Cdc42-2ritC model result; values with Cdc42-GTP levels below this line are postulated to lead to inviability. Gray regions in patch radius plot indicate conditions where sGAP is not strongly affected by exo/endocytosis and no stable patch forms.

Despite the opposing membrane flow, all three simulated Cdc42-ritC formed an active patch of enhanced Cdc42-GTP and depleted of sGAP, though very marginally for Cdc42-3ritC (Figure 3A, Movies S2-S4). The profiles of total Cdc42 as well as Cdc42-GTP reproduce the experimental trends, showing decreasing concentrations with decreasing Cdc42 mobilities (Figure 3B, 2F). Thus, GEF-mediated positive feedback in concert with flow-displacement of sGAP oppose the diluting effect of membrane flow on Cdc42, even for Cdc42 mobilities within the depletion region of Figure 2B. However, reduction of Cdc42 mobility results in reduced overall Cdc42-GTP activation and a broadened patch, as shown in more detail in a systematic parameter scan of dissociation and diffusion constants (Figure 3C).

### Reducing GAP activity restores viability to cells with slow Cdc42

The simulation results of Figure 3 show that in regions corresponding to Cdc42-3ritC mobility the number of Cdc42-GTP particles is very low. This observation leads to the hypothesis that a minimal level of Cdc42 activity, at a “survival” threshold above the Cdc42-3ritC parameters, is necessary for cell viability (Figure 3C). Given the poor viability of Cdc42-2ritC, this survival threshold may overlap with the Cdc42-2ritC measured in diploid cells. We thus tested if a reduction in GAP activity in the model might bring Cdc42-3ritC above the survival threshold. Indeed, reducing the amount of sGAP particles by half or removing sGAP completely enlarges the viability region (Figure S5) and brings Cdc42-3ritC simulated cells at or above the survival threshold (Figure 4A).

**Figure 4.**
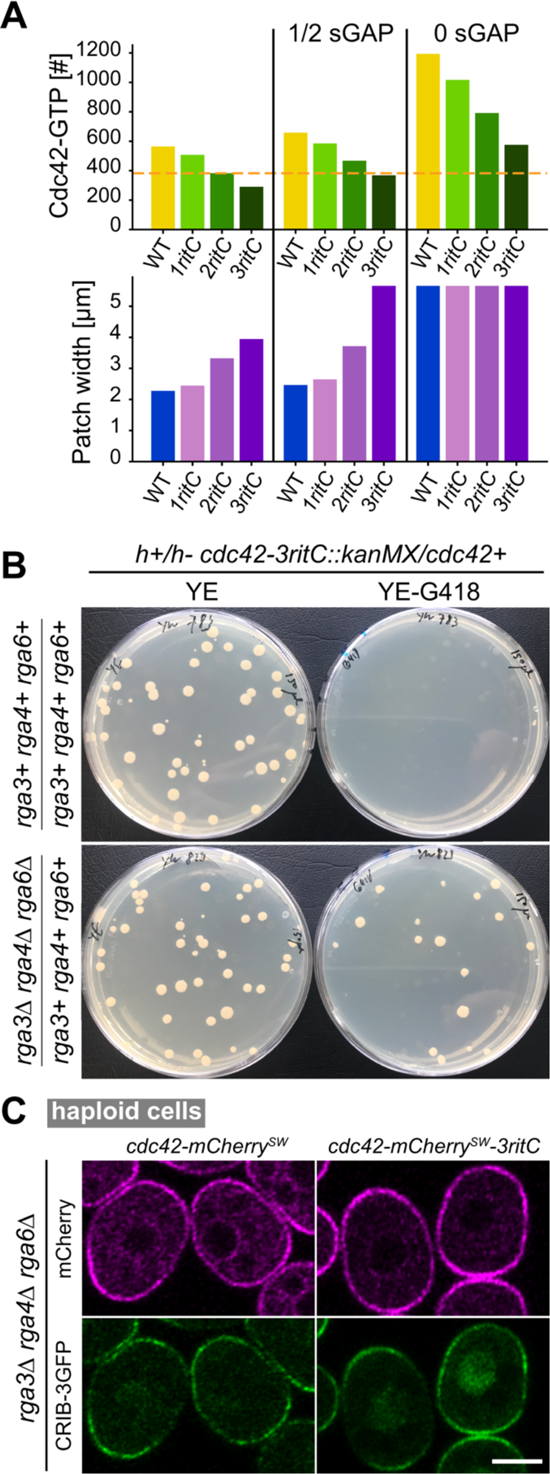
Deletion of Cdc42 GAPs restores viability to strains containing low-mobility Cdc42. **A.** Model predictions for total amount of Cdc42-GTP and width of the polarity patch in reference conditions and upon reduction of sGAP. **B.** Random spore analysis from heterozygous *cdc42-mCherry^SW^-XritC / cdc42+* diploids, which were otherwise WT (top) or contained heterozygous deletions for each of the three Cdc42 GAPs (bottom). Growth on rich non-selective (YE) and *cdc42-mCherry^SW^-XritC*-selective (YE-G418) plates is shown. **C.** Airyscan2 images of Cdc42-mCherry^SW^ variants and CRIB-3GFP (marking Cdc42-GTP) in a haploid strain lacking all Cdc42 GAPs. Scale bar is 5µm.

To test experimentally if reducing Cdc42-GTP hydrolysis by GAPs restores viability to *cdc42-3ritC* cells, we constructed the *cdc42-3ritC* allele in a diploid strain that also carried heterozygous deletions of *rga3*, *rga4* and *rga6*. Sporulation of the strain would in principle permit recovery of haploid progeny carrying any combination of the *cdc42-3ritC* allele with *rga3Δ*, *rga4Δ* and/or *rga6Δ*. Remarkably, whereas sporulation of the *cdc42-3ritC/+* diploid did not produce G418-positive colonies (as also shown in the tetrad dissection, Figure 2G), sporulation of the *cdc42-3ritC/+ rga3Δ/+ rga4Δ/+ rga6Δ/+* diploid produced G418-positive colonies, indicating recovery of viable *cdc42-3ritC* haploid cells (Figure 4B). These cells were also mutant for *rga3Δ*, *rga4Δ* and/or *rga6Δ*, and we readily recovered haploid triple *rga3Δ rga4Δ rga6Δ* mutant cells that carried *cdc42-3ritC* as sole *cdc42* allele (Figure 4C). These cells were very abnormally shaped, similar to the *rga3Δ rga4Δ rga6Δ* triple mutants carrying a WT *cdc42* allele (28), and had strong CRIB signal at the plasma membrane, showing that reduction of Cdc42-GTP hydrolysis by removal of Cdc42 GAPs restores Cdc42-3ritC activity. Thus, deletion of Cdc42 GAPs restores viability to a low-mobility Cdc42 allele.

## Discussion

Reaction-diffusion models have been very successfully used to explain pattern formation on membrane surfaces. However, membranes are not inert surfaces, but may be actively remodeled by pattern-forming proteins at its surface, an aspect not generally captured in reaction-diffusion models. For instance, a local patch of Cdc42 GTPase activity promotes polarized secretion, inevitably leading to membrane flows away from the patch. Here, we developed a particle-based model to explicitly probe how membrane flows contribute to Cdc42 GTPase symmetry breaking. Our findings, supported by experimental validation, demonstrate that, in the presence of membrane flow, differential dynamics of fast Cdc42 and slow inhibitor are critical for Cdc42 pattern formation. Thus, membrane flows enable protein dispersion that both constrains the membrane affinity of polarity factors and allows long-range displacement of inhibitors, implementing a transport-driven lateral inhibition mechanism as in classic activator-inhibitor pattern formation (31).

The model demonstrates how membrane flows can be beneficial for polarization, by promoting the exclusion of a slow GAP. As a patch of Cdc42-GTP forms, local exocytosis promotes sGAP depletion, forming a zone permissive for Cdc42 activation. The slowness of GAP dynamics further helps to pin the location of the established activation zone. In absence of membrane trafficking, sGAP is not depleted from the forming Cdc42-GTP patch and inactivates Cdc42. The simulations best reproduce experimental Cdc42-GTP patch formation in cells expressing optoGAP as sole Cdc42 GAP (22). OptoGAP distributes homogeneously at the plasma membrane and turns off Cdc42 in the dark, but tightly binds the plasma membrane upon illumination, allowing its concomitant depletion from zones of polarized secretion and emergence of a Cdc42-GTP patch, similar to the emergence of the Cdc42-GTP patch in simulations. The endogenous GAPs Rga4 and Rga6 localize to cell sides and are excluded from zones of active secretion. We had previously shown that Rga4 localization is largely governed by membrane flow dynamics (22). The localization of Rga6, which is slightly less depleted from cell poles than Rga4, and its dependence on polarized secretion are also consistent with patterning by membrane flows (25), though this has not been directly tested. The contributions of these GAPs to Cdc42 patch formation are likely more complex as additional signals, such as lipid specificity or the cell cycle, also modulate their localization (32, 33). Fission yeast cells also express another Cdc42 GAP, Rga3, that colocalizes with the Cdc42-GTP patch (28). Fine dissection of GAP localization and development of alleles that do not deplete from cell poles will be required to probe how much GAP depletion by membrane flows reinforces Cdc42-GTP patch formation in the natural in vivo context.

We show, both in silico and experimentally, that sufficiently fast dynamics (fast membrane unbinding rates and/or fast lateral diffusion) of Cdc42 is required for efficient polarization. The loss of polarization upon decreased Cdc42 mobility can be understood in the following way: Arrival of secretory vesicles at the patch causes local displacement and dilution of Cdc42, leading to widening and weakening of the patch. Consequently, secretory vesicle delivery is less polarized, reducing the net membrane flow. sGAP is thus less efficiently displaced, causing stronger overlap between sGAP and Cdc42-GTP and consequent Cdc42 inactivation. By contrast, fast dynamics allow Cdc42 to rapidly repopulate the zones of secretion, alleviating the effects of the membrane turnover. We conclude that establishment and maintenance of a polarized patch requires its components to turnover faster than the surface on which it is formed.

Additional flows may exist to aid cell polarization. For instance, Cdc42 is itself delivered to cell poles on secretory vesicles (9, 19, 34). Thus, even though the concentration of Cdc42 in secretory vesicles was measured to be lower than that at the plasma membrane and dilute the polarity patch (18, 19), secretory vesicles do not arrive “naked” with respect to Cdc42. While vesicle-based influx may not play a decisive role for the highly mobile WT Cdc42 (17), it may be important for slower molecules. In previous work, we had shown that anchoring Cdc42 at the plasma membrane with a transmembrane anchor (Cdc42-TM), which cannot unbind the membrane but is delivered to cell poles on secretory vesicles, led to a viable allele, in contrast to the Cdc42-3ritC we present here (9). Transmembrane proteins could be less perturbed by membrane flows in comparison to peripheral membrane proteins, due to interaction with structures that remain stationary with respect to lipid flow (accounted for by model parameter γ > 1). This suggests that membrane unbinding and rebinding is not the only mechanism that can efficiently repopulate Cdc42 at the poles.

We note that, with the Cdc42 rates we measured, the model predicts a somewhat stronger depletion of Cdc42-1/2ritC than we observe experimentally. Some of this discrepancy may stem experimentally from the selective pressure for viability of haploid cells. For instance, survivor *cdc42-2ritC* cells, at the edge of viability, may be selected to be in the high range of the Cdc42-2ritC mobility distribution. It is also likely that some aspects of Cdc42 polarization are not fully captured by the positive feedback and membrane flow simulations and that there are other positive inputs in the system, such as those played by microtubule-delivered factors (35, 36).

Membrane flows are likely to play important roles in cell patterning across organisms (23). For instance, whereas budding yeast cells do not exhibit Cdc42 GAPs depleted from the polarity patch, the formation of the septin ring, which forms a structural base of the bud neck and is proposed to associate with Cdc42 GAPs, involves central depletion by exocytosis (37). A similar setup may be at play in pollen tubes, where the Rho GTPase inhibitor REN4 is displaced to the edges of pollen tips by the growth process (38). While cell polarity activators and inhibitors may vary across organisms, polarized secretion-induced membrane flows are likely to be intrinsic to polarization mechanisms across cell types.

## Materials and Methods

### Mathematical Modeling Methods and FRAP analysis

#### Computational model of Cdc42 polarization with explicit GAPs and membrane flow by exocytosis and endocytosis

##### Simulation Domain and Parameters

We simulated proteins as a system of particles on a surface representing the plasma membrane that migrate due to both reaction-diffusion and membrane delivery/removal by exocytosis/endocytosis. We focus on a single tip of the fission yeast cell, represented as a sphere with radius 1.8 μm. The particles are either restricted to the surface of this sphere or present in a cytoplasmic pool. Positions of particles in this cytoplasmic pool are not kept track of since we assume cytoplasmic diffusion is fast. The rate constants in the simulation were informed by FRAP experiments (this work), based on prior studies, or estimated as explained below (Tables S1, S2).

##### Diffusion

Particle diffusion on the spherical domain surface, with diffusion coefficient *D*, is implemented with Brownian dynamics as in (22). Briefly, the magnitude of the displacement due to diffusion over time Δt is calculated as √2DΔtξ(t) independently in both directions in the plane tangent to the sphere at the original particle location, and ξ(t) is a random Gaussian distributed variable with zero mean and unit variance. After calculating the displacement in the tangent plane, the positions of each particle are then projected down to the sphere surface while keeping the distance conserved (39).

##### Particle reactions

Reactions of diffusing particle species in the simulation include membrane binding and unbinding, unimolecular reactions (indicated with rate constants symbols *k*, r in Figure 1A), and bimolecular reactions on the membrane (indicated by λ in Figure 1A). When particles on the sphere undergo an unbinding event, they are removed from the sphere surface and placed in an internal pool representing the cytoplasm. Particles in this internal pool are placed back on the surface anywhere with uniform probability according to the association rate constants. The probability of a reaction occurring in time Δt for a given particle or pair of particles is equal to *P_react_* = 1 − exp (−rΔt), where r is the reaction rate, with the exception of membrane binding which was handled with the Gillespie algorithm (where the time until the next reaction is a continuous variable and the reaction that occurs is chosen randomly according to the sum of all reaction rates that can occur through this method (40)). Bimolecular reactions in the model include discrete sGAP hydrolysis, Scd1/2/Cdc42-GTP trimer complex formation, and promotion of Cdc42 to its active form by either Scd1/2 or Scd1/2/Cdc42-GTP. These reactions depend on the existence of two particles and most are indicated by reactions that have a λ variable indicating their rate (Figure 1A). Bimolecular reactions with rate λ are implemented using the Doi method where two species within a microscopic distance, *ρ*, react with rate *λ* (29, 41–43). Using this method, *ρ* and *λ* together define the macroscopic rate constant. For Scd1/Scd2/Cdc42-GTP formation via cytoplasmic Scd1/2 binding to membrane-bound Cdc42-GTP, we calculate the rate by the rate constant *k_ST_* instead of the Doi method (since the positions of cytoplasmic particles are not defined in our model). When particles form a complex (reaction between Scd1/2 and Cdc42-GTP particles, both bound to the membrane), the complex is placed at the midpoint between the prior positions of the two reactants. When complexes separate, the separated particles are placed at distance *σ,* slightly larger than ρ. Hydrolysis and activation reactions change the type of Cdc42 particle with the sGAP and Scd1/Scd2 complex positions unchanged.

##### Assumptions of particle reaction scheme

The rate constant of hydrolysis *k_hydro_* represents the combined effect of spontaneous hydrolysis and hydrolysis by the GAP protein Rga3, which is known to colocalize with Cdc42-GTP to the tips of *S. pombe* cells (28). We assume that *k_hydro_* is uniform in space. (2) Hydrolysis by slow GAP (sGAP) represents hydrolysis by Rga4/6 which localizes to the sides of *S. pombe* cells in part due to membrane flows (22). For simplicity, and without loss of generality, we assume sGAP has low mobility with a diffusion coefficient of 10^-3^ μm^2^/s and no membrane binding and unbinding, causing it to be influenced by membrane flows and redistribute away from patches of active Cdc42 (locations of exo/endocytosis). (3) Reactions involving Scd1/2 represent a minimal version of the positive feedback required for symmetry breaking (11) that approximates the sum of separate contributions of Scd1, Gef1, and Ras1 (2, 44, 45).

##### Membrane flows

Particle movement by exocytosis and endocytosis was implemented as in prior work (22). Briefly, we assume that lipid traffic exocytosis and endocytosis near cell tips create a flow of lipids and of all associated proteins included in the model along the plasma membrane. We do not incorporate a full hydrodynamic model, assuming the instantaneous displacement of proteins around the site of the vesicle event (18, 46). This assumption is supported by the appearance of a flat plasma membrane in electron micrographs of fission yeast cell at cell sides and tips, except at sites of vesicle traffic or around eisosomes (47–49), similar to S. cerevisiae (48, 50). This observation indicates the presence of a membrane tension, which, combined with turgor pressure, can flatten the vesicle membrane delivered by exocytosis (51, 52), causing lateral flow. Similarly, flow of lipids towards endocytic vesicles must occur over a sufficiently wide area, preventing membrane strains larger than a few % that would otherwise lead to formation of membrane pores (53).

Our model incorporates three coarse-grained hydrodynamic parameters. (1) Parameter α describes the mobile fraction of lipids in the plasma membrane; where α = 1 indicates that the entire membrane is able to flow. We used α = 0.5. (2) Parameter γ measures the ratio of lipid to membrane-bound protein velocity, which could be larger than unity due to interactions of proteins with structures that remain stationary or less mobile with respect to lipid flow, such as in the cell wall or in the cytoplasm. We used γ = 1. (3) Parameter *R_cutoff_* is a cutoff distance to the flow induced by individual exocytosis and endocytosis events, accounting for slow equilibration of membrane tension over long distances (54). We used *R_cutoff_* = 2 µm. Protein depletion by membrane flow is robust with respect to the precise values of α, γ, and *R_cutoff_* (22).

Lipids on the surface at arc length distance s from the center of an exocytosis event are assumed to be pushed away radially by arc length distance Δr(s). This displacement is calculated by assuming that the difference between the spherical cap surface area corresponding to mobile lipids measured at the final and initial arc length positions of the particle from the center of exocytosis equals the surface area of the exocytotic vesicle *A_exo_*. The displacement is only applied to particles up to an upper arc length distance, s < *R_cutoff_*. For an endocytosis event, particles within the cutoff distance *R_cutoff_* are moved radially towards the endocytic point (or exactly at the endocytic point if they happen to be sufficiently close to it). The displacement calculation is similar to exocytosis and considers the surface area of the endocytic vesicle, *A_endo_*. We thus assumed that the particles of our model do not get internalized by endocytosis.

##### Exo/endocytosis distributions

We assume that both exocytosis and endocytosis events occur with constant overall rates, directed towards sites of membrane-bound Cdc42-GTP. Specifically, that each Cdc42-GTP enhances the probability of an exo/endocytosis event occurring near it according to a Gaussian distribution as function of arc-length distance to the Cdc42-GTP particle, with standard deviation *W_exo_* or *W_exo_*, respectively. The probability distribution of an event occurring anywhere on the simulation domain is the sum of Gaussian distributions centered on each Cdc42-GTP particle position (Figure 1B). We tuned the values of *W_exo_* or *W_exo_* to reproduce the experimentally-measured exocytosis and endocytosis profiles around the cell tip as determined from Exo70-GFP and Fim1-mCherry (22), given the experimentally-measured profile of Cdc42-GTP measured by CRIB-GFP (9). This implies 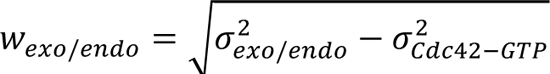, where σ*_exo/end_^2^* is the variance of the experimental exocytosis/endocytosis profile and σ*_cdc42-GTP_^2^*is the variance of the experimental CRIB profile as function of arc-length distance from the cell tip. This tuning gives *W_endo_* = 1.51 µm. Since the experimental exocytosis profile is slightly narrower than the Cdc42-GTP profile, *W_exo_* is set to a small value (0.01 μm). At reference parameters, the model generates exo/endocytosis distributions that are in good agreement to experimentally measured profiles (Figure S3).

##### Time Ste

The diffusion of particles on the spherical surface was implemented using Brownian dynamics with a fixed timestep of 10^-3^ s. An upper bound for this timestep of 7.35 × 10*^47^*s was calculated from the root mean squared displacement equation for diffusion on 2D surfaces, ρ ≈ √2DΔt, where D is the largest diffusion constant used in the simulation (0.17 μm^2^/s). We truncated this upper bound to determine a timestep of 10^-3^ s sufficient for our reaction parameters. Additionally, for the largest λ we consider a bimolecular reaction occurs on the first step after entering the reaction radius, ρ, slightly less than 1% of the time.

##### Simulations with varying Cdc42 mobility

In simulations of Cdc42-xritC constructs, when lowering the Cdc42-GDP diffusion coefficient and membrane dissociation rate constant, we also lowered the corresponding values of Cdc42-GTP and Scd1/2/Cdc42-GTP so that Cdc42-GDP would always have greater than or equal mobility compared to Cdc42-GTP. Specifically, the three following equations hold throughout all simulations *D_T_ = min{D_D_, D_T,WT_}, r_T_ = min {r_D_, r_T,WT_},* and *D_ST_* = min{*D_T_, D_ST,WT_*} where subscript WT indicate WT values (Table S1). In simulations of Cdc42-1ritC, Cdc42-2ritC, and Cdc42-3ritC constructs, we also assume the number of Cdc42 on the membrane is approximately the same as in WT cells, though our results are robust with respect to changes in this number. To achieve this, and similar to the WT case (Table S1), the membrane binding rate of Cdc42-GDP was set to (r*_’_*/2 + r*_8_*) where r*_’_* is the dissociation rate constant of Cdc42-xritC-GTP and r*_8_* is the dissociation rate of Cdc42-xritC-GDP.

##### Diploid simulations

For diploid simulations, we doubled the number of each species in the system and assigned half of these the parameters WT parameters for Cdc42 unbinding and diffusion while the other half was assigned dissociation and diffusion constants for a different reference case of either Cdc42-WT, Cdc42-1ritC, Cdc42-2ritC, or Cdc42-3ritC.

##### Quantification of particle concentration profiles

Particle concentration profiles (e.g. Figure 1E) were measured by calculating the arclength distance from the centroid associated with the largest cluster of Cdc42-GTP. Cdc42-GTP particles were placed in the same cluster if their arclength distance was smaller than 0.3 μm. The instantaneous centroid was calculated by averaging the positions of Cdc42-GTP in this largest cluster, and then projecting back to the sphere surface. The centroid used for the profiles was calculated from an average over five frames of the instantaneous largest centroid. The arclength of the particles on the sphere were measured from this centroid and then placed into histograms for each particle type. The values in this histogram were divided by the area of the strip on the sphere surface to determine the surface density for each particle. Profiles were averaged over at least 500 s.

##### Quantification of polarization patch width

Patch widths were calculated based on the sGAP profile. Specifically, by determining the arclength from the Cdc42-GTP centroid value at which the sGAP concentration first drops below 90% of the concentration sGAP has far from the patch centroid. The far field concentration was determined by averaging the sGAP concentration over 5.05 μm to 5.55 μm arclength from the patch centroid. The arclength of the first drop below 90% was determined by starting from an arclength of 5.05 μm and moving towards 0 μm, averaging concentration over a moving window of 0.1 μm. Cases where the concentration of sGAP did not drop below 90% of the far field value were marked as unpolarized (Figures 3C, S5B) or the patch width was set equal to half the circumference of the sphere, 5.65 μm (Figure 4A).

### Determination of membrane diffusion and dissociation rates by FRAP

Effective diffusion coefficients D and membrane dissociation constants *k_+--_*for WT and Cdc42 mutants were estimated by fitting the experimental recovery profiles to a 1D model as described in prior work (22).

#### FRAP of cell sides

Experimental fluorescence intensity profiles were acquired after photobleaching of a rectangular region and imaging on the top surface of *S. pombe* cells (Figure S2A). Since Cdc42 is primarily inactive at cell sides (9), we assume the fit for D and *k_off_* corresponds D*_8_* and r*_8_* for Cdc42-GDP. This fit for two parameters relies on D and *k_off_* contributing differently to the recovery profile, with diffusion providing smoothing of profile over time and association/dissociation contributing to uniform recovery (in the limit of fast cytoplasmic diffusion). The values of these two parameters were further narrowed down by performing FRAP of different bleach widths, which have varying relative recovery contributions of diffusion and association/dissociation.

We corrected for photobleaching by multiplying all intensities by *e^kt^* after background subtraction, where *k* is the exponential decay constant of the average intensity of a neighboring non-bleached cell. From these photobleach-corrected images we determined a 1D intensity profile *I*(*x*,*t*) (where the x-dimension corresponds to the long axis of the cell) by defining a rectangular region of interest along the long axis of a bleached cell with width ∼2 μm, excluding the curved region close to the cell tips, and calculated the average intensity over time along this line at points spaced 0.0516 μm (1 pixel) apart using ImageJ’s getProfile function.

Each FRAP fit to *I*(*x*,*t*) results to individual best fit points in diffusion and dissociation constant space (Figures 2B, S2E). The dashed contour lines in Figures 2B, S2E were determined by averaging together the normalized R^2^ measure of fits for each FRAP curve and finding the contour line that traces out 96% of the maximum. The maximum value of this average normalized R^2^ was taken as the best fit value for as the diffusion and dissociation constants.

#### Model with diffusion and association/dissociation used for fitting

We modelled the evolution of the concentration profile of a membrane-associated protein, *c_m_*(*x*, t), along a 1D line with reflecting boundaries at positions *x_a_*, *x*_b*=*_, defining the finite size of a membrane domain. We assumed *c_m_*(*x*, t) obeys the following equations, which include both the effects of diffusion and membrane binding/unbinding:

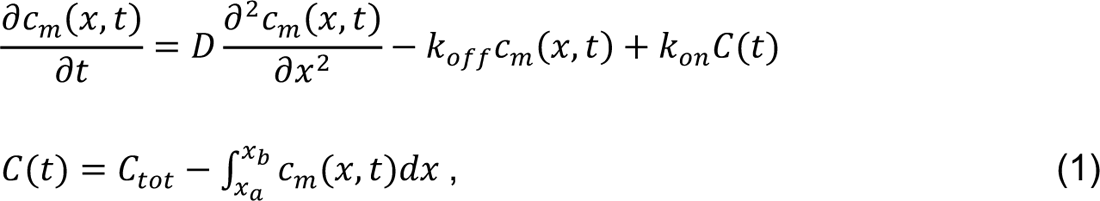

where *c_m_* is concentration on the membrane, *C*_tot_ is the number of proteins in the cytoplasm, *C_tot_* is the total number of proteins, and *k_off_*, *k_on_*are the membrane dissociation and association rate constants. We do not write a separate diffusion equation for the cytoplasm, assuming that cytoplasmic diffusion is sufficiently fast. We also specifically consider the limit where most of the proteins are on the membrane: *C*(t) ≪ *C_tot_*, (equivalently, the limit of sufficiently large *k_on_*). The Green’s function describing the probability for a single protein being at position *x* at time t, given that it was at position *x_>_* at time t = 0 is

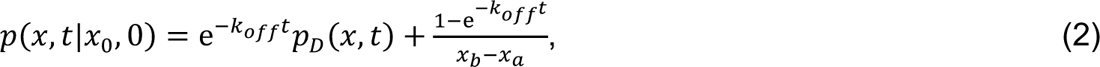

where *p_D_* (*x*, t) accounts for the probability of the particle position due to diffusion alone:

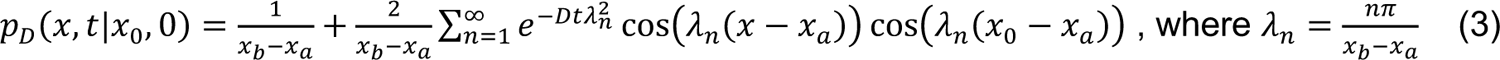

#### Fitting to experimental I(x,t) profiles

Assuming *c_m_*(*x*, t) is proportional to the experimental *I*(*x*, t), we solve for *c_m_*(*x*, t) given an initial condition *c_m_*(*x*, 0) that is proportional to the intensity profile imaged immediately after the photobleaching, *I*(*x*, 0). We then fit the model to both *x* and *t* after photobleaching simultaneously, using SciPy’s optimization curve fitting to get a single best fit value for D and *k_+--_* as described below. The sum in Eq. (3) was carried up to *n* = 500. The intensity profile is divided into three regions: left, middle, and right. The intensity in the middle region, defined to be between *x_*_* and *x_”_* where *x_a_* ≤ *x_d_* ≤ *x_e_* ≤ *x_b_*, is determined by the intensity profile *I*_i_*^0^*immediately after photobleaching, evaluated at pixels *i* of the projected profile, corresponding to positions *x_E_*. The middle region is therefore treated as a sum of *n* = Δ*x*/(*x_e_* – *x_d_*) separate integrals, where Δ*x* is the pixel size. The lengths of the left and right flanking regions are both equal to *l_F_*, with *l_F_* = *x_d_*− *x_a_* = *x*_b*=*_ − *x_e_*. We average the intensity of the pre-bleached cell over the middle region and set this as the initial intensity of the two flanking regions, *I*_F_*^0^*. The value of *l*_F_ determines the uniform intensity at long times, *I_B_*, according to mass conservation:

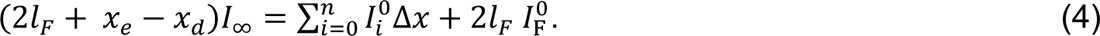

The long time intensity *I_B_*, or equivalently *l_F_*, is determined via the fitting procedure. When the *l_F_* value drifted towards 0 or large numbers, it was restricted to be in the range 2-10 µm. The equation describing the concentration or intensity as a function of position *x_E_* corresponding to pixel *i*, and time is found by analytical integration, *I* _model_(*x*, t) = ∫_xa_^xb^ *I*(*x^I^*, 0)*p*(*x*, t|*x*′, 0)p*x*′ (22) with the initial intensity *I*(*x*, 0) at *x_E_* < *x* < *x_EJ?_* given by 0.5(*I^>^* + *I^>^*). An example of a fit over time to the experimental FRAP data for one cell with Cdc42-WT is shown in Figure S2.

#### Fit to half-tip FRAP

We obtained estimates for diffusion coefficient and dissociation constants for Cdc42-GTP in WT cells by bleaching half of the cell tip (9, 55). We fitted the same model of Equations (1)-(4) to the photobleached-corrected intensity profile along a contour drawn along the cell periphery of a medial confocal section (Figure S2C,D). Assuming that Cdc42-GTP dominates the recovery, we use the fit for D and *k_+--_* to estimate D*_’_* and r*_’_* (Figure S2E). We only fitted the short time recovery (< 5 s) where Cdc42-GTP diffusion is expected to contribute to broadening of the boundary between the bleached and unbleached regions while dissociation to uniform recovery, prior to Cdc42-GDP diffusion to the tip from the cell sides. We note that this is only a rough estimate of the effective Cdc42-GTP mobility, given the differences between the 1D model and 3D geometry and the fact that the tip FRAP does not become homogenous at long times as assumed by the 1D model.

### Experimental methods

#### Media and growth conditions

For microscopy experiments, cells were grown to exponential phase at 25°C in Edinburgh minimal medium (EMM) supplemented with amino acids as required, except diploids which were grown in complete YE medium to repress sporulation. Tetrad dissections and dilution series were grown on YE plates at 25°C, unless otherwise indicated. Random spore analysis was done by digestion of sporulated diploids with glusulase, growth on YE plates at 25°C and replication on YE plates containing appropriate antibiotics.

#### Strain construction

Strain genotype is given in Table S3. Integrative plasmids carrying the *cdc42-3ritC* and *cdc42-2ritC* alleles were constructed as follows. Plasmid pSM1442 (pFA6a-3′UTR(*cdc42)-cdc42-mCherry^SW^-1ritC*-Term-kanMX; (9)) was digested with NheI-XmaI to remove the 1ritC fragment. The 3ritC fragment, amplified with primers osm7870 and osm7871 from pAV601, and the 2ritC fragment, amplified with primers osm7870 and osm7872 from pAV602 (22), were cloned by InFusion cloning, yielding plasmids pSM2934 (pFA6a-3′UTR(*cdc42)-*cdc42-mCherry*^SW^*-3ritC-Term-kanMX) and pSM2935 (pFA6a-3′UTR(*cdc42)-*cdc42-mCherry*^SW^*-2ritC-Term-kanMX), respectively. These plasmids were linearized with SalI and integrated in diploid strains constructed using the trans-complementation of *ade6-M210* and *ade6-M216* alleles. In each case, the diploids were selected on EMM, and propagated on YE plates, before transformation and selection on YE-G418 plates.

The *cdc42-2ritC* and *cdc42-3ritC* alleles in otherwise WT diploid strains were obtained by transformation of linearized pSM2934 and pSM2935 in a strain obtained by crossing haploids *h+ ura4-D18 ade6-M210* (YSM4072) and *h-ura4+ ade6-M216* (YSM4073), yielding strains YSM4074 and YSM4075, respectively (Figure 2G). The same alleles in a strain carrying the CRIB marker were obtained by transformation in a diploid obtained by crossing haploid strains *h-ura4-D18 ade6-M210 leu1-32:pshk1:CRIB-3GFP:ura4+:leu1+* (YSM4076) with *h+ ura4-D18 ade6-M216 leu1-32* (YSM1182), yielding strains YSM4077 and YSM4078, respectively. The *cdc42-3ritC* allele in a strain carrying deletion of genes encoding Cdc42 GAPs was obtained by transformation of linearized pSM2934 in a diploid obtained by crossing *h+ ura4-ade6-M216 leu1-rga4Δ::natMX rga3Δ::hphMX* (YSM4079) and *h-ura4-D18 ade6-M210 leu1-32:pshk1:CRIB-3GFP:ura4+:leu1+ rga6Δ::bleMX* (YSM4080), yielding strain YSM4081.

The diploid strain carrying the *cdc42-1ritC* allele was obtained by crossing a derivative of the *cdc42-1ritC* strain described in (9), *h+ ura4-D18 ade6-M216 leu1-32 cdc42-mCherrySW-1ritC:kanMX* (YSM4082), with YSM4076, yielding strain YSM4083.

Haploid *cdc42-2ritC* strains were obtained through sporulation and tetrad dissection of diploids: YSM4084 from sporulation of YSM4075 (Figure 2H); YSM4085 from sporulation of YSM4078 (Figure 2E); and a haploid *cdc42-3ritC* strain *rga3Δ rga4Δ rga6Δ* deletion (YSM4086) from sporulation of YSM4081. As this strain did not contain the CRIB transgene, this was re-integrated yielding strain YSM4087 (*rga3Δ rga4Δ rga6Δ*) (Figure 3C). Diagnostic PCRs distinguishing WT from *cdc42-mCherry-3ritC* alleles were used to further to verify the Cdc42 allele present in this strain.

### Microscopy

Cells were imaged with Plan-Apochromat 63×/1.40 oil differential interference contrast objective on an LSM980 system equipped with an airyscan2 detector and acquired by the ZEN Blue software (Zeiss). Imaging was set in SR mode with frame bidirectional scanning. Laser power was kept at <0.6%, with pixel time around 13.2 µs, and frame size of 634×634. All other settings were optimized as recommended by the ZEN Blue software. Images in Figures 2E and 4C were acquired with 4x averaging to obtain smoother distributions of CRIB and Cdc42. FRAP at cell middle was performed on cells placed on an EMM-2% agarose pad by finding the focal plane closest to the coverslip and defining a rectangular region of interest of defined dimension (1 to 6µm wide) in the middle of the visible signal (see Figure S2A). FRAP at cell poles was performed on the medial focal plane using a rectangle covering half of the cell pole (see Figure S2C). Two snapshots were taken before bleaching, which was done using a single pass of the bleach zone at 100% laser power. Recovery was imaged for up to 5 min at intervals adapted to the speed of recovery of each protein. All Airyscan images were processed in ZEN Blue software.

### Image analysis

Profiles of Cdc42 and CRIB at the cell periphery were acquired in FIJI from 6 µm-long (7µm-long for diploid cells), 5 pixel-wide segmented lines starting at the cell pole with higher CRIB signal (2 profiles per cell) and averaged. To estimate the amount of fluorescence signal captured in the cortical profiles, we assumed that any CRIB-GFP signal in cortical profiles at the middle of interphase WT cells represents cytosolic signal. We found that this corresponds to about half of the background cytosolic fluorescence measured in a small region devoid of vacuoles. We thus removed cytosolic signal from cortical profiles by subtracting 0.5 x the cytosolic fluorescence measured in a small region devoid of vacuoles. The CRIB profiles plotted in Figures 2D and 2F are averages (≥ 24 profiles for averaged images in 2F; ≥ 84 profiles for non-averaged images in 2D). To compare the tip to side distribution of Cdc42, we subtracted the average value at distance 5-6µm from the cell pole from the Cdc42 profiles.

## Acknowledgements

We thank Laetitia Michon for help with molecular biology. Work in SGM’s lab is supported by Swiss National Science Foundation grants 310030_191990 and 310030_207909, and European Research Council Advanced grant 101019630 (SexYeast). DV and DMR were supported by NIH grant R35GM136372. Portions of this research were conducted on Lehigh University’s Research Computing infrastructure partially supported by NSF Award 2019035 and on Expanse Cluster at SDSC through allocation TG-MCB180021 from the Advanced Cyberinfrastructure Coordination Ecosystem: Services & Support (ACCESS) program, which is supported by NSF grants #2138259, #2138286, #2138307, #2137603, and #2138296.

## Supporting Information for

**Figure S1.**
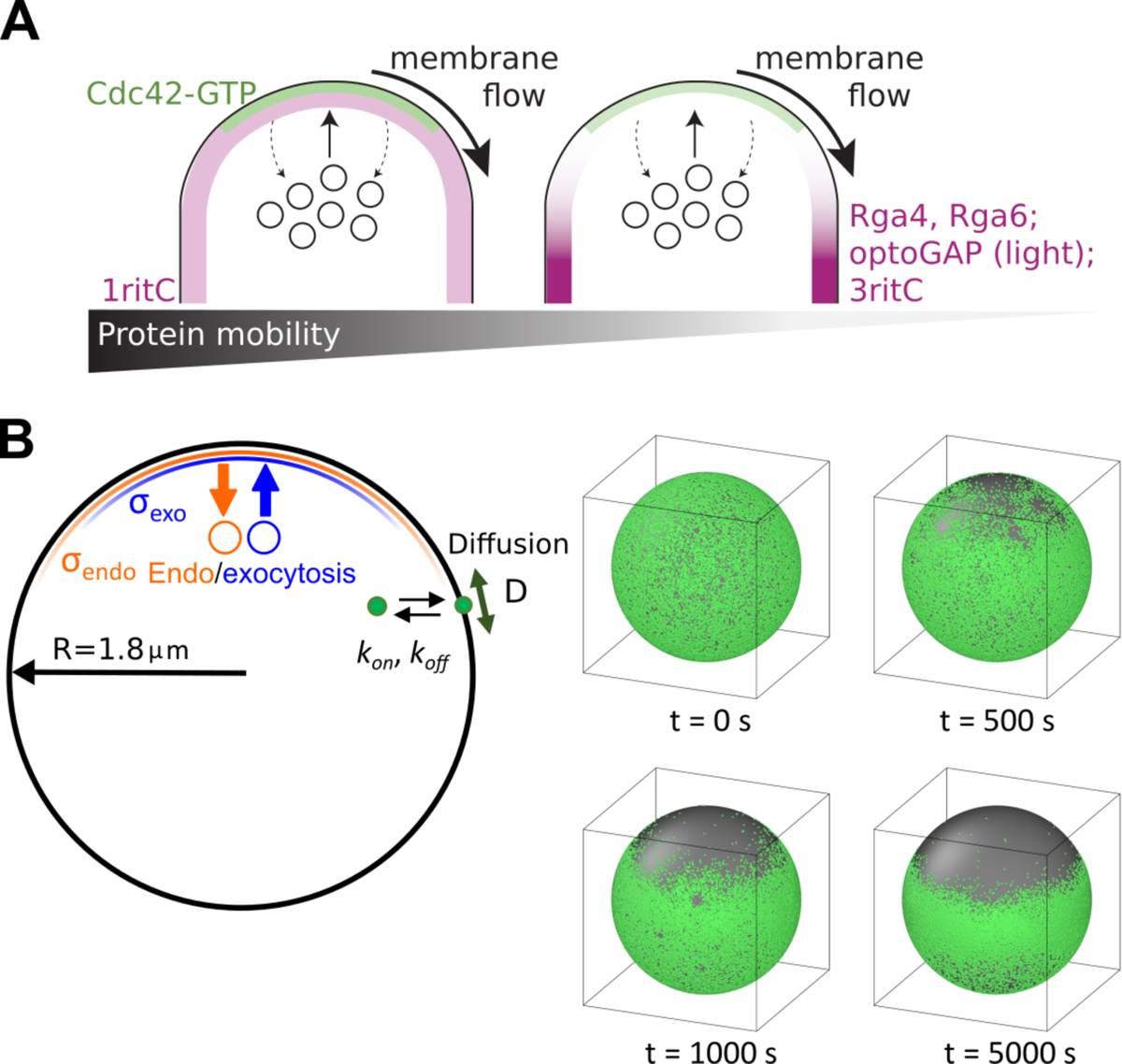
In plane membrane flow due to localized secretion and broader endocytosis can lead to peripheral membrane protein depletion. (A) Schematics showing that high mobility proteins (Cdc42-GTP, 1ritC) are able to keep up with membrane flows due to exocytosis and endocytosis and remain at the tip of growing *S. pombe* cells. Low mobility proteins (Rga4/6, optoGAP which dimerizes under light, and 3ritC) are displaced by the membrane flow. (B) Computational model of membrane flow, reproduced from (1), simulated as stochastic, area-conserving exocytosis and endocytosis events centered at a point of spherical domain, with inert particles undergoing diffusion and binding/unbinding. Model shows depletion of initially uniform particle distribution away from the region where exocytosis and endocytosis are occurring, for sufficiently small diffusion coefficients and unbinding rates. The depletion phenomenon occurs when the exocytosis region is narrower than the endocytosis even when the overall rates of membrane secretion and internalization are equal to each other.

**Figure S2.**
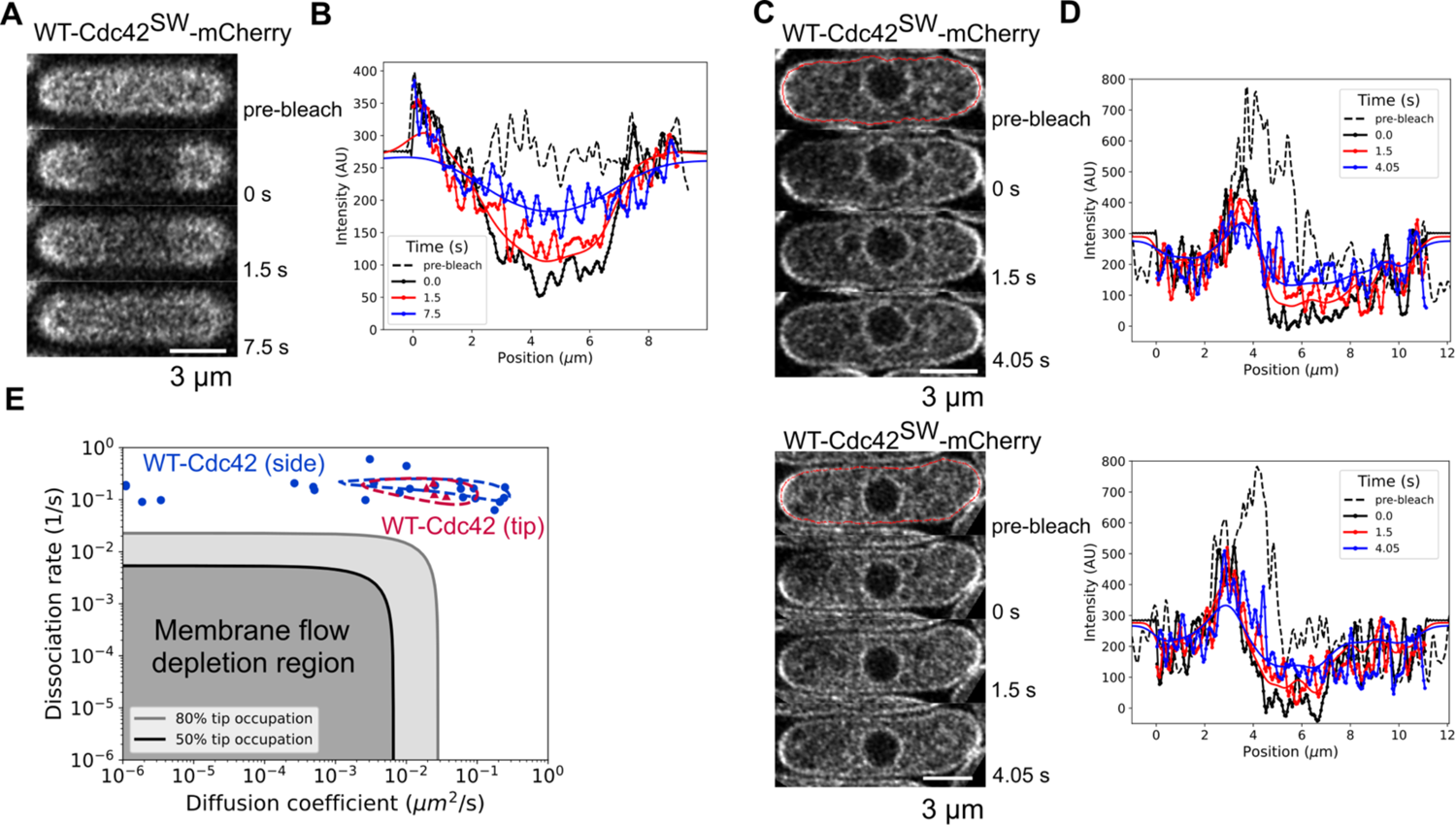
FRAP fits to extract diffusion coefficients and membrane dissociation rates. (A) Photobleaching of a rectangular region along the cells sides. Confocal section at the cell surface for WT Cdc42-mCherry^SW^. Similar FRAP experiments were performed for cells in Figure 2B. (B) Recovery traces over time and fit (solid lines) for cell in panel A. The fitted intensity was measured by projecting the intensity within a box of size the cell width perpendicularly to the long axis of the cell. (C) Two examples of half-tip bleach of WT Cdc42-mCherry^SW^, imaged along a confocal section through the cell middle. (D) Recovery traces over time along with model fit (solid lines) for cells in panel C. The intensity was measured over a strip of width four pixels (1 pixel = 0.0516 μm) along a cell contour shown as red line in panel C. (E) Best fit parameters for membrane unbinding and diffusion for both side and tip recovery as in Figure 2B. Dashed line regions are drawn based on an average across all cell recoveries. Cdc42-GDP diffusion and dissociation constants are comparable to ∼0.2 µm^2^/s and ∼0.03 /s measured using a different FRAP method in (5).

**Figure S3.**
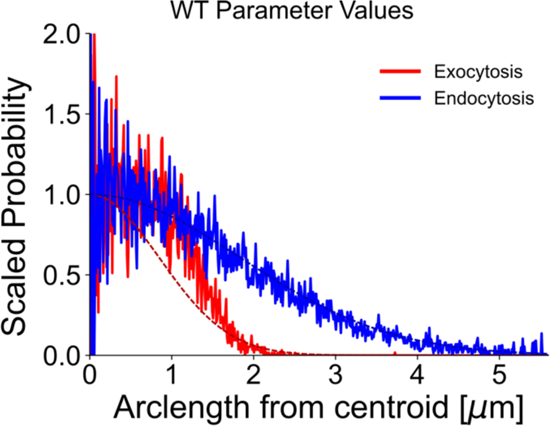
Steady state distribution of exocytosis and endocytosis in WT simulations of Figure 1E (solid lines) versus the desired distribution (dashed lines) in (1). Average was calculated similarly to concentration profiles over 5000 s at steady state.

**Figure S4.**
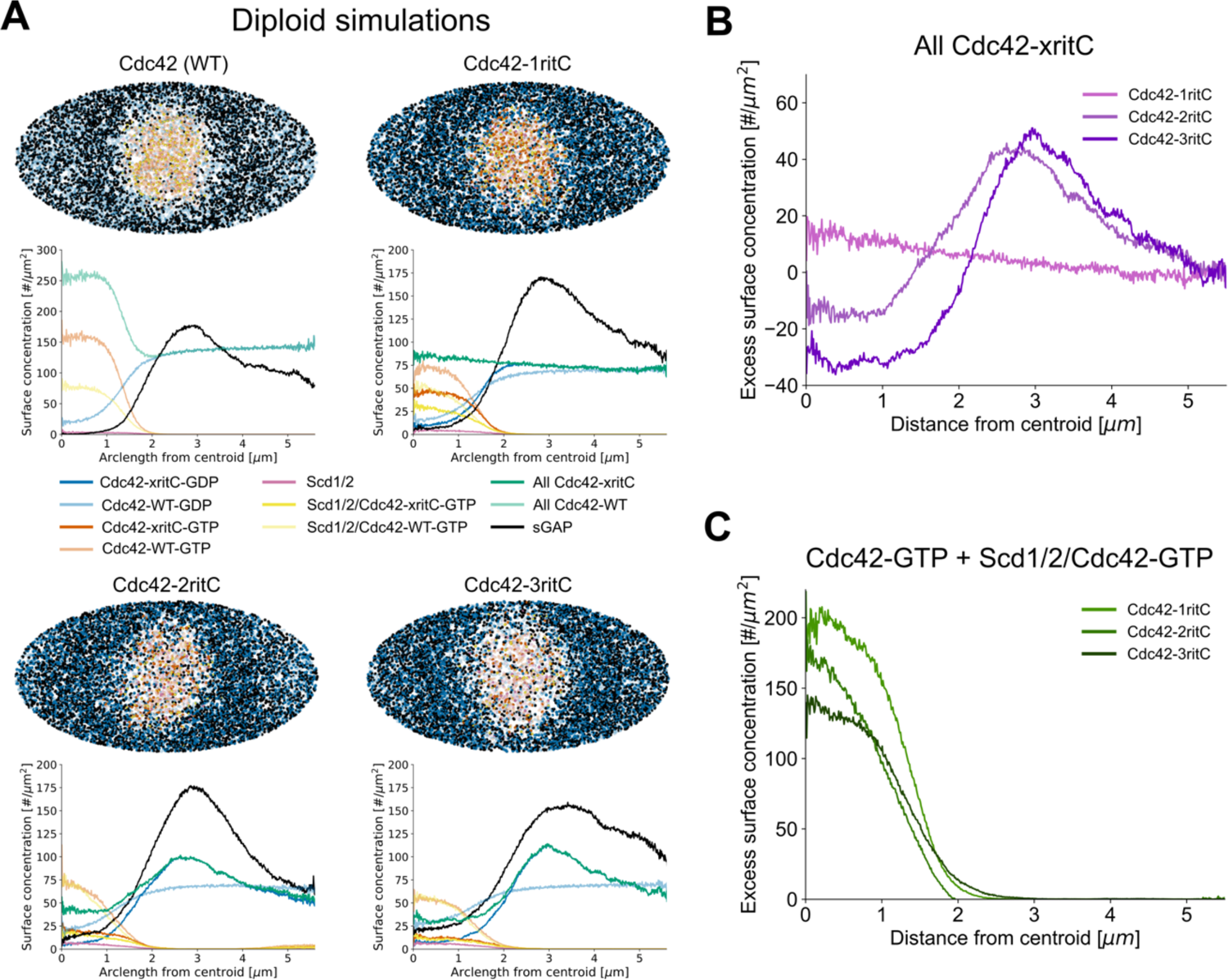
Simulated polarization for diploid cell parameters. (A) Steady state snapshots and concentration profiles from diploid simulations after steady state for WT or where one copy (half of Cdc42) has lower mobility corresponding to either Cdc42-1ritC, Cdc42-2ritC, or Cdc42-3ritC. WT Cdc42 associated particles shown in faded colors. Profiles are averaged over time as in Figure 1. Snapshots were taken after 600 s. (B) Side normalized Cdc42 concentration of only the mutant Cdc42 component. (C) Side normalized Cdc42-GTP concentration for both mutant and WT components.

**Figure S5.**
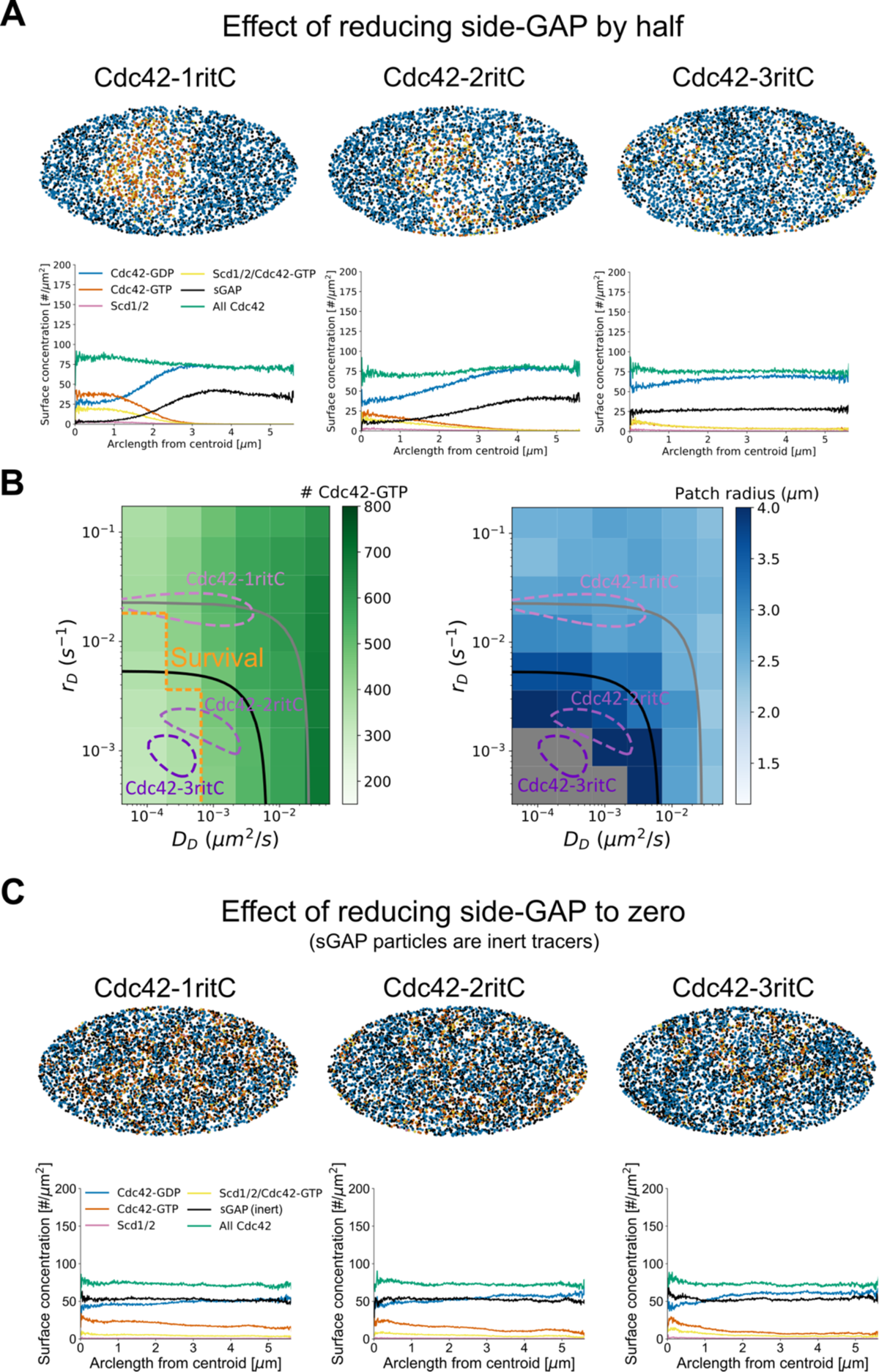
Simulations with reduction of sGAP particle numbers. (A) Steady state snapshots and concentration profiles for simulations where the number of sGAP has been reduced by ½ compared to Figure 1E. The simulation with Cdc42-ritC shows small transient Cdc42-GTP clusters that are picked up in the concentration profile. However, these small clusters are mobile over the simulation time and do not lead to a depletion of sGAP around them. Snapshots were taken after 14000 s. (B) Parameter scan for amount of Cdc42-GTP and the patch width as in Figure 3E, for ½ sGAP simulations. The survival threshold line shrinks compared to Figure 3E. (C) Steady state snapshots and concentration profiles for simulations where the number of sGAP has been reduced to 0. In these simulations passive tracer sGAP particles were added to detect stable polarization. All cases show transient Cdc42-GTP clusters that are picked up in the concentration profile. These small clusters are mobile over the simulation time and do not lead to a depletion of inert tracer sGAP around them. Snapshots were taken at 15000 s.

**Table S1.**
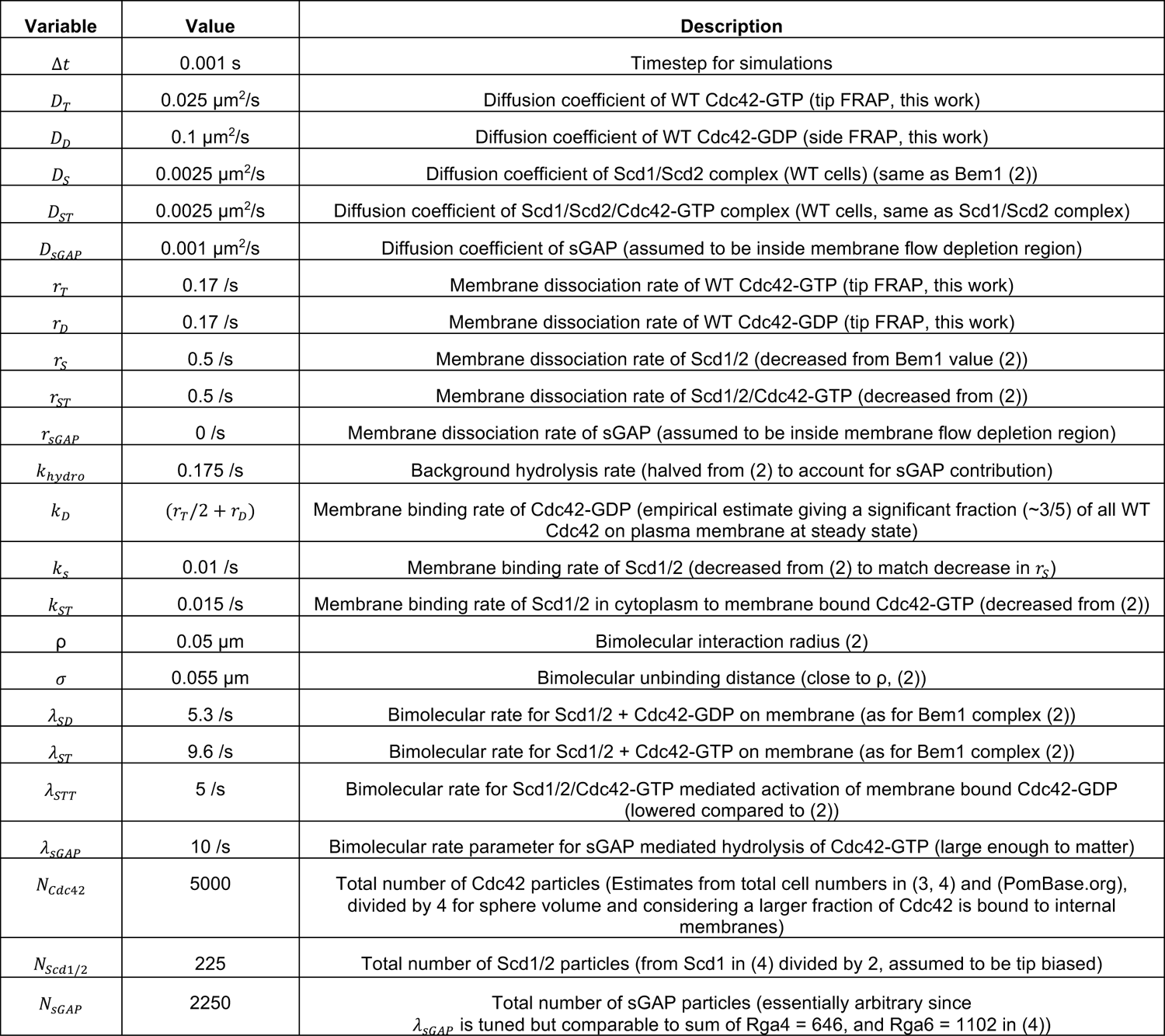
Model parameter values for particle reactions.

**Table S2.** Model parameter values for membrane flows from (1) unless otherwise indicated.

|  |  |  |
| --- | --- | --- |
| $A_{exo}$ | $3.14 \times 10^{-2} \mu\text{m}^2$ | Surface area of individual exocytotic vesicle |
| $A_{endo}$ | $6.42 \times 10^{-3} \mu\text{m}^2$ | Surface area of individual endocytotic vesicle |
| $r_{exo}$ | 0.68 /s | Rate of exocytosis events per tip in growing *S. pombe* |
| $r_{endo}$ | 2.44 /s | Rate of endocytosis events per tip in growing *S. pombe* |
| $w_{exo}$ | 0.01 $\mu\text{m}$ | Variance of blurring Gaussian for exocytosis (determined based on profiles from (5) and (1)) |
| $w_{endo}$ | 1.51 $\mu\text{m}$ | Variance of blurring Gaussian for endocytosis (determined based on profiles from (5) and (1)) |
| $\alpha$ | 0.5 | Fraction of mobile component in membrane |
| $\gamma$ | 1.0 | Coupling between the proteins and the flowing membrane |
| $R_{cutoff}$ | 2.0 $\mu\text{m}$ | Cutoff for effect of exo/endocytosis |

**Table S3.** Strain list.

| Strain number | Genotype | Reference |
| --- | --- | --- |
| YSM3138 | h+ leu1-32 cdc42-mCherry<sup>SW</sup>:Term:kanMX ura4-294:pshk1:CRIB-3GFP:ura4+ | (5) |
| YSM2468 | h+ leu1-32 cdc42-mCherry<sup>SW</sup>-1ritC:kanMX ura4-294:pshk1:CRIB-3GFP:ura4+ | (5) |
| YVV4074 | h+/h- ura4-/ura4+ ade6-M210/ade6-M216 cdc42-mCherry<sup>SW</sup>-3ritC:Term:kanMX/cdc42+ | This work |
| YSM4075 | h+/h- ura4-/ura4+ ade6-M210/ade6-M216 cdc42-mCherry<sup>SW</sup>-2ritC:Term:kanMX/cdc42+ | This work |
| YSM4077 | h+/h- ura4-D18 ade6-M210/ade6-M216 leu1-32:pshk1:CRIB-3GFP:ura4+:leu1+/leu1+ cdc42-mCherry<sup>SW</sup>-3ritC:Term:kanMX/cdc42+ | This work |
| YSM4078 | h+/h- ura4-D18 ade6-M210/ade6-M216 leu1-32:pshk1:CRIB-3GFP:ura4+:leu1+/leu1+ cdc42-mCherry<sup>SW</sup>-2ritC:Term:kanMX/cdc42+ | This work |
| YSM4081 | h+/h- ura4- ade6-M210/ade6-M216 leu1-32:pshk1:CRIB-3GFP:ura4+:leu1+/leu1- rga6Δ::bleMX/rga6+ rga4Δ::natMX/rga4+ rga3Δ::hphMX/rga3+ cdc42-mCherry<sup>SW</sup>-3ritC:kanMX | This work |
| YSM4083 | h+/h- ura4-D18 ade6-M210/ade6-M216 leu1-32/leu1-32:pshk1:CRIB-3GFP:ura4+:leu1+ cdc42-mCherry<sup>SW</sup>-1ritC:kanMX/cdc42+ | This work |
| YSM4084 | ura4? ade6- cdc42-mCherry<sup>SW</sup>-2ritC:Term:kanMX | This work |
| YSM4085 | ura4-D18 ade6- leu1-32:pshk1:CRIB-3GFP:ura4+:leu1+ cdc42-mCherry<sup>SW</sup>-2ritC:Term:kanMX | This work |
| YSM4087 | ade6- rga6Δ::bleMX rga4Δ::natMX rga3Δ::hphMX cdc42-mCherry<sup>SW</sup>-3ritC:kanMX leu1-32:pshk1:CRIB-3GFP:ura4+:leu1+ | This work |

**Movie S1.** Simulation with WT cell parameters (same as Figure 1E) from 0 to 500 s. 2D Mollweide projection of particles on a sphere.

**Movie S2.** Simulation with Cdc42-1ritC haploid cell parameters (same as Figure 3A) from 0 to 1500 s. 2D Mollweide projection of particles on a sphere.

**Movie S3.** Simulation with Cdc42-2ritC haploid cell parameters (same as Figure 3A) from 0 to 5000 s. 2D Mollweide projection of particles on a sphere.

**Movie S4.** Simulation with Cdc42-3ritC haploid cell parameters (same as Figure 3A) from 0 to 9000 s. 2D Mollweide projection of particles on a sphere.

